# Humans, but not songbirds, produce spontaneously isochronous song through the lifespan

**DOI:** 10.1101/2025.06.02.657467

**Authors:** Mila Bertolo, Michael W. Weiss, Isabelle Peretz, Jon T. Sakata

**Author notes:** These authors contributed equally (Isabelle Peretz, Jon T. Sakata).

## Abstract

Isochrony – as in the regular beat of a metronome - is cross-culturally ubiquitous in music. Is this ubiquity due to a widespread biological inclination for acoustic communication having isochronous structure? If so, it should be present in lesser-studied vocal music, in the absence of musical training, and in comparable non-human species’ vocalizations. We quantified isochrony in an untrained expression of musicality: improvised songs from non-musician adults and children. We also tested for isochrony in songs from zebra finches, a bird that learns complex songs. We analyzed improvised songs from children (n = 38, 3-10 years old) and adult non-musicians (n = 15, 24-82 years old), and songs from juvenile and adult zebra finches (n= 77, ∼50 and 120 days post-hatch, respectively). Isochrony was expressed in non-musicians’ improvised songs, and in children’s improvised songs to a comparable degree. In contrast, both juvenile and adult zebra finches produced songs with less isochrony than chance. And although zebra finches learn sequences and durations of their songs’ elements, we found no evidence for the learning of isochrony. These data show that spontaneous isochrony in learned vocalizations appears differently across these two species. We propose that species variation in chorusing behaviors may explain why.

## Introduction

Rhythm is one of the basic building blocks of human musicality^1^. In its simplest form, the use of a regular pulse (like the ticking of a clock or metronome) - referred to as *isochrony* - is extremely widespread among music traditions^2,3^. Musicians from Mali, Bulgaria, and Germany all finger-tap to reliably mimic small integer ratio rhythms, where durations of adjacent notes have small integer ratio relationships, such as 1:1 and 1:2; they also show a tendency to simplify complex rhythms (e.g. to 1:1, isochrony)^4^. Such cross-cultural ubiquity suggests the behavior has some shared, biological basis. Indeed, the ability to clap in time to a perceived isochronous pulse in music has an identifiable polygenetic basis in humans^5^. The presence of isochrony in the vocalizations of distantly-related species - non-human primates^6–9^, songbirds^10,11^, frogs^12^, and pinnipeds^13–15^ - further indicate some biological basis to this behavior. To date, however, we have little data concerning whether isochrony is expressed in an ancient and universal musical form - singing - and whether there are comparable expressions of isochrony in non-human species capable of combinatorial, learned vocalizations (as opposed to simple, repeated, innate ones). Here, we present a developmental and comparative analysis of isochrony in human improvised songs, and spontaneous song from a non-human animal that is capable of learned, complex songs: the zebra finch songbird.

Isochrony production appears to be widespread and consistent in human percussive musical behaviors. Musicianship is not a prerequisite: non-musicians’ accuracy in tapping synchronously to isochronous stimuli is on par with that of amateur musicians^16^. When musicians and non-musicians from 15 countries were tasked with reproducing an auditory rhythm by tapping along to it, their “errors” systematically favored isochrony even when it was not originally present in the stimulus^3^. Percussive isochrony emerges early in human development; by 2-4 years old, children’s manual tapping behaviors spontaneously show regular rates^17^, and they show full-body rhythmic movements in response to music (though inconsistently synchronized to it)^18^. By 6-7 years old, the ability to synchronize to an isochronous stimulus is almost at adult-level accuracy^19^, and this accuracy is well-preserved through ageing (reviewed in^20^).

If isochrony is as fundamental to human musicality as the above findings suggest, we should also see this rhythm in untrained vocal song. Singing is a universal^21^ and spontaneous behavior in humans, and its perception is associated with specific neural networks^22^. Young children create new tunes and adapt familiar ones^23,24^, and this practice of real-time improvisation is found across many of the world’s musical traditions^25^. Even those with little musical training can improvise songs, and those songs remarkably show pitch use that is consistent with the conventions of Western music (in Western participants)^26^. Improvisational singing, furthermore, is a methodologically useful tool in that it can be intuitively used by people with little to no formal musical training, through the lifespan. Previous findings of isochrony in corpora that include both instrumental and vocal music^2^ suggest that isochrony is likely also present in song, but such corpora are the product of explicit musical training. Years of explicit musical training is an exception rather than a norm, making the study of musicians a steep under-representation of human musicality; instead, the study of non-musicians’ improvised song – in the absence of explicit training or mimicry – can better characterize how spontaneously isochrony may arise in human musicality. To our knowledge, isochrony in improvised singing has never been examined. We expect that non-musicians’ improvised songs should also show isochrony in line with their percussive musical behaviors. Similarly, this trait should emerge around the age of 6 or 7 in children’s improvised songs.

This method of sung improvisation, which by design imposes very little constraints on participants, also allows for more direct cross-species comparisons. Very few non-human species “improvise”; many species’ vocalizations consist of innate calls that are repeated at consistent and stereotyped intervals^12,27^ creating, by definition, isochronous sequences. By contrast, vocalizations that combine and sequence many different sound elements do not necessarily result in an isochronous rhythm; we know much less about whether isochrony is also the prevailing rhythmic organization in the combinatorial vocalizations of non-human animals.

Songbirds like the zebra finch are ideal for investigating isochrony in learned vocalizations. Zebra finches learn their song by memorizing the sound of and practicing how to imitate a tutor’s song, and mechanisms underlying the acquisition, performance, and perception of these learned songs have been extensively studied^28^. Zebra finches learn the sequencing and timing of vocal elements (“syllables”) in their songs^28,29^, affording them a level of temporal control that in principle could be used to give their complex vocalizations rhythmic structure similar to human song. Timing is ecologically relevant for these songbirds; they coordinate their song with body and head movements^30,31^, and show inter-individual coordinated timing^32^, but notably they do not sing synchronously. Zebra finches are also able to perceptually discriminate between isochronous and nonisochronous stimuli with some degree of tempo flexibility^11^.

Here, we provide a developmental and comparative analysis of isochrony in humans and songbirds, focusing on spontaneous/improvised songs in each species. We tested for the presence of isochrony in the improvised songs of children and adult non-musicians, and in the spontaneous songs of juvenile and adult zebra finches. While two previous studies have reported isochrony in adult zebra finch song^10,33^, no study to date has examined developmental changes in isochrony, analyzed the contribution of learning to isochrony, and directly compared isochrony in zebra finch song to isochrony in human vocal music. This comparative study of isochrony will provide a more detailed account of the degree of similarity between human and songbird vocal rhythms are, which will help evaluate candidate theories for the origins of rhythm, including the vocal learning hypothesis for beat perception and synchronization^34^, the gradual audiomotor evolution hypothesis^35^, and the origins of rhythm production in group dancing^36^ or other group behaviors^37,38^.

## Methods

We tested for the presence of isochrony in the following three datasets:

### 1. Human, cross-sectional

We analyzed recordings of spontaneous songs from adults (n = 15, ages 24-82 years old) and children (n = 38, ages 3-10 years old). Procedures for adult and children song recordings were approved by the Comité d’éthique de la recherche en éducation et en psychologie (CEREP), University of Montreal.

#### Adults

15 non-musician adults were recorded as part of a prior study^26^. To summarise, these participants (*M* = 58.6 years, *SD* = 21.3 years; 10 female, 5 male) resided in Quebec, Canada. Those who requested information were told that the task involved singing. They did not have specific music deficits associated with congenital amusia using the Montreal Protocol for Identification of Amusia^39^, and were considered “non-musicians” in that they had 5 or fewer years of music training. Participants provided informed consent and were paid for their participation.

The participants were instructed to “make up a new melody” for each improvisation trial, that should not be a repetition of a melody they had heard before, or a repetition of a melody they produced in a previous trial. They were told to improvise using the syllable “da” instead of using words, and to sing for as long as felt appropriate. Each participant completed 28 improvisations; 8 in response to verbal prompts (lullaby, dance, healing, and love songs; 2 happy, 2 sad songs), 16 prompted with short melodic stems (3-4 notes), 2 prompted with longer melodic stems (10 notes), and 2 periods of free improvisation. Participants were tested while seated in a sound-attenuating booth, where melodic prompts were presented to them via headphones (DT-770 Pro, Beyerdynamic, Inc.) through Audition (Adobe, Inc.). Verbal prompts were communicated to them by the experimenter, who was outside the booth and able to communicate with the participant through the participant’s headphones. Their improvisations were recorded via a Neumann TLM-103 microphone.

#### Children

38 children (M = 6.3 years old, range 3.4 - 10.4; 22 female, 16 male) were recruited using social media to diffuse an advertisement for an online study on children’s improvised singing, with the incentive that participating families could be entered in a raffle to earn a $50 gift card to the bookstore of their choice. By following the hyperlink in the ad, parents were directed to an online study hosted on https://brams.org/, run on jspsych^40^. After a parent provided informed consent and answered demographic questions, the game provided audio instructions for musical games for the child. An onscreen character (“Bunny” in English, “Lapinou” in French) narrated instructions for the child to sing back songs that they played (these warm-up trials were not included in analysis), and then to make up a song *“without any words, like “da da da”, and try to sing something totally new that you’ve never heard before!”*. This prompt for made-up songs was repeated 6 times, and after each of these trials, a text prompt addressed to the parent asked whether the song was truly improvised, or whether it was familiar to them; only songs that the parent marked as improvised were retained for analysis. Children produced an average of 2.4 songs (range 1 - 5, with an average of 83.9 notes (SD = 78.3) per trial. The total study ran for approximately 10 minutes. Like the adults in our study, the children had less than 5 years of music training (musical instrument lessons: *m* = 0.8, *SD* = 1.4, range = 0 - 5; or voice lessons *m* = 0.2, *SD* = 0.9, range 0 - 5).

## 2. Zebra finch colony population

61 male zebra finches were semi-naturally raised in a breeding colony at McGill University. These birds were raised by their parents and learnt their song from their father. They remained with their parents until they were ∼60 dph and then socially housed in same-sex group cages. All these birds were recorded when they were developmentally mature (i.e., >120 dph). 40 of the birds in the dataset were raised by 21 different fathers (on average, each father raised and tutored 1.9 pupils; range 1 - 5 pupils per tutor). Four of these 61 birds were also represented in the longitudinal dataset (with only their adult recordings included in this population dataset).

### 3. Zebra finch, longitudinal

The 16 birds in this analysis were raised by both parents, and were able to physically and acoustically interact with other birds. Only male zebra finches learn to produce songs, so only males were analyzed here. Male juveniles were individually housed in a sound-attenuating chamber for a few days for song recording (see below) starting at 44-51 days post-hatch (dph). The birds were then housed with other males until they reached adulthood (∼120 dph), at which point they were moved back into the sound-attenuating chamber for song recording for a few days, and then returned to a group cage.

All birds were housed on a 14 L:10 D light cycle and provided food and water ad libitum. For all bird recordings, zebra finches were recorded whilst housed individually in a sound-attenuating chamber (TRA Acoustics, Ontario, Canada) via an omnidirectional microphone (Countryman Associates, Inc, Menlo Park, CA). A continuously recording system (Sound Analysis Pro (SAP) 2011) was set to save audio as triggered by the detection of sound (digitized at 44.1 kHz). Like humans, zebra finches produce songs even without audiences^41^, and all songs recorded for this study were taken while the birds were housed individually; all these songs are therefore spontaneous (“undirected”) song. All procedures were approved by the McGill University Animal Care and Use Committee in accordance with the guidelines of the Canadian Council on Animal Care.

## Data Processing

For all datasets, onsets of notes (in humans) and “syllables” (in birdsong) were semi-automatically timestamped. Human recording audio files were processed in TONY^42^ (Fig. 1). The automatically generated note onsets and offsets were manually checked, and corrected where necessary (e.g., erroneous boundaries placed by the algorithm). For each zebra finch, 10 recordings per bird were semi-automatically annotated using custom software in MATLAB (`evsonganaly`). Adult zebra finch songs are highly stereotyped, with the timing and sequencing of syllables being highly consistent across renditions; as such, the rhythm of a bird’s song can be readily captured with few song examples. Similar to many published studies (e.g.,^29^), segmentation parameters were determined per bird (i.e., amplitude threshold above which one would consider there to be a sound; the signal generally has to be above this threshold for at least 20ms to mark a syllable, and below this threshold for at least 5ms to mark a gap). Audio files can contain both songs and calls; all segmented sounds were manually annotated in order to retain only song syllables for subsequent analysis. All vocalizations within songs (i.e. introductory notes and song syllables) were annotated for analysis. We used a threshold of 1 second of silence as the delineator between separate bouts.

**Fig. 1.**
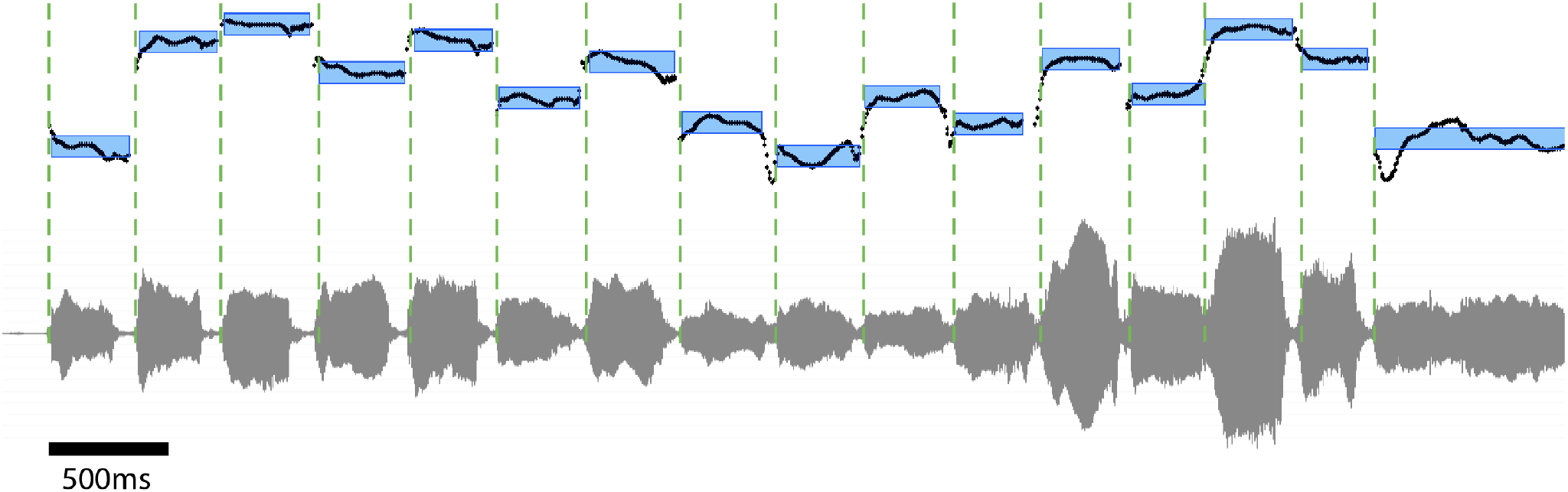
Example waveform (grey, bottom) and mean frequency (black, top) of a child’s improvised song. Blue boxes delineate TONY automatic segmentation (see Methods). Visually, the intervals between the onsets (green dashed lines) of adjacent blue bars are roughly equivalent, indicating the isochronous organization of this song. We used the dyadic interval ratio to quantify isochrony in the song recordings of human children and adults, and zebra finch juveniles and adults.

## Analyses

We computed inter-onset intervals (*IOI* s) from the recordings’ timestamped onsets of notes and syllables. We quantified the presence of isochrony using the ***dyadic interval ratio*** (as in^6,10,14,43^): for any given dyad i.e., two adjacent inter-onset intervals, this is the ratio resulting from the duration of the first IOI divided by the sum of the whole dyad’s IOIs: 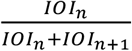. For a recording of a ticking clock, for example, where all IOIs are identical, one would only observe the ratio 0.5. For a recording of any signal that is more temporally variable than a ticking clock, one would observe a distribution of dyadic interval ratios between the bounds [0, 1], where the density of ratios around 0.5 may be taken as indicative of how isochronous this signal is. As in prior research^6^, we accepted a small window of ratio values around 0.5 as indicative of isochrony (ratio values in the window [0.444, 0.555]; henceforth referred to as the *isochrony window*), and a small range of values symmetrical around this window as being “off” isochrony (windows [0.400-0.444] and [0.555-0.600]). When comparing the density of ratio values that fall “on” isochrony relative to “off”, we normalized the number of ratios by dividing this count by the width of the window they are observed in (also consistent with^6,9^). This is a characterization of isochronous organization of a recording’s notes/syllables’ IOIs, so we note that this is different than a study of a “beat” which could refer to a perceived, latent isochronous pulse; a beat can be perceived from a stimulus that does not have acoustic events at every beat, and has extra acoustic events in between beats.

The above analysis approach that compares the density of ratios in the isochrony window to the density of ratios outside of it tests the observed data against an implicit, relatively weak null hypothesis of randomly generated interval lengths^44^. We sought to additionally test a stricter null hypothesis that accounts for there being production constraints and biases on interval lengths. In order to quantify whether observed isochrony is higher than would be expected by chance, we simulated 50 datasets in which the order of IOIs were scrambled for each participant, and distributions of dyadic interval ratios re-computed for these data. These simulated datasets are therefore biologically plausible in that they retain each participant/bird’s propensity to produce IOIs of certain durations, but reduces any rhythmic structure to only what may be observed by chance from randomly sequenced vocalizations.

We tested for the presence of isochrony in each population (children, adults, zebra finch juveniles, zebra finch adults) first by applying paired t-tests to determine whether the normalized count of dyadic interval ratios was greater inside the isochrony window than “off” this window. In order to test whether each population showed more isochrony than chance, we applied paired t-tests comparing each participant’s number of ratios in the isochrony window in actual data vs the average incidence in their scrambled data.

To test whether, in either species, there were developmental differences in the incidence of isochrony, we summarised each participant’s data into a proportion of how many observed ratios fell in the isochrony window. Because proportions are a measure bounded [0,1] and therefore best modeled with a beta distribution, we fit beta regressions for each species (package *glmmTMB*) predicting this outcome variable *proportion*, using a fixed effect for age group (juvenile/children vs adults), and random intercepts for ID in the longitudinal bird data (where data is grouped by ID). We fit another beta regression to test whether human children’s ages correlated with the proportion of ratios that fell in the isochrony window (fixed effect for children’s ages). Zebra finches learn many temporal aspects of their song, including duration of syllables and gaps, and the sequencing of syllables. We therefore tested whether the degree of isochrony in their song also appeared to be learned. We tested this by running a beta regression modelling the proportion of a pupil’s ratios in the isochrony window, as predicted by a fixed effect for the tutor’s proportion of ratios in isochrony window, with a random intercept for tutor ID (as some tutors had more than one pupil).

We additionally computed descriptive statistics for the human data to contextualise how similar our recordings of improvised song are to previous studies of song and instrumental music. This included average speed (the inverse of IOI length) per group, and coefficient of variation (see SI). We tested whether adults’ lullaby vs dance songs differed in speed by fitting a mixed model testing mean IOI as predicted from the fixed effect prompt type (lullaby or dance song) and random effect for participant ID. We used t-tests to test whether children and adults’ songs differed in speed; and a Pearson correlation to test whether children’s ages correlated with speed. These analyses indicated that our samples of improvised songs bore similarities to prior studies of musical behaviours, in that children had faster spontaneous production rates than adults, and that improvised lullabies were slower than improvised dance songs (see SI for details). These samples of improvised song therefore appear to be ecologically valid samples that are comparable with musical behaviours reported elsewhere, on these basic dimensions.

## Results

### 1. Adult non-musicians’ improvised songs display isochrony

Adult non-musicians’ improvised songs showed significantly more dyadic interval ratios in the isochrony window than the off window (*Figure 2A*; *t*(14) = 6.37, *p* < .001). Additionally, the degree of isochrony in these songs was greater than one would observe by chance (*Figure 2B*; *t*(14) = 5.579 *p* < .001).

**Fig. 2.**
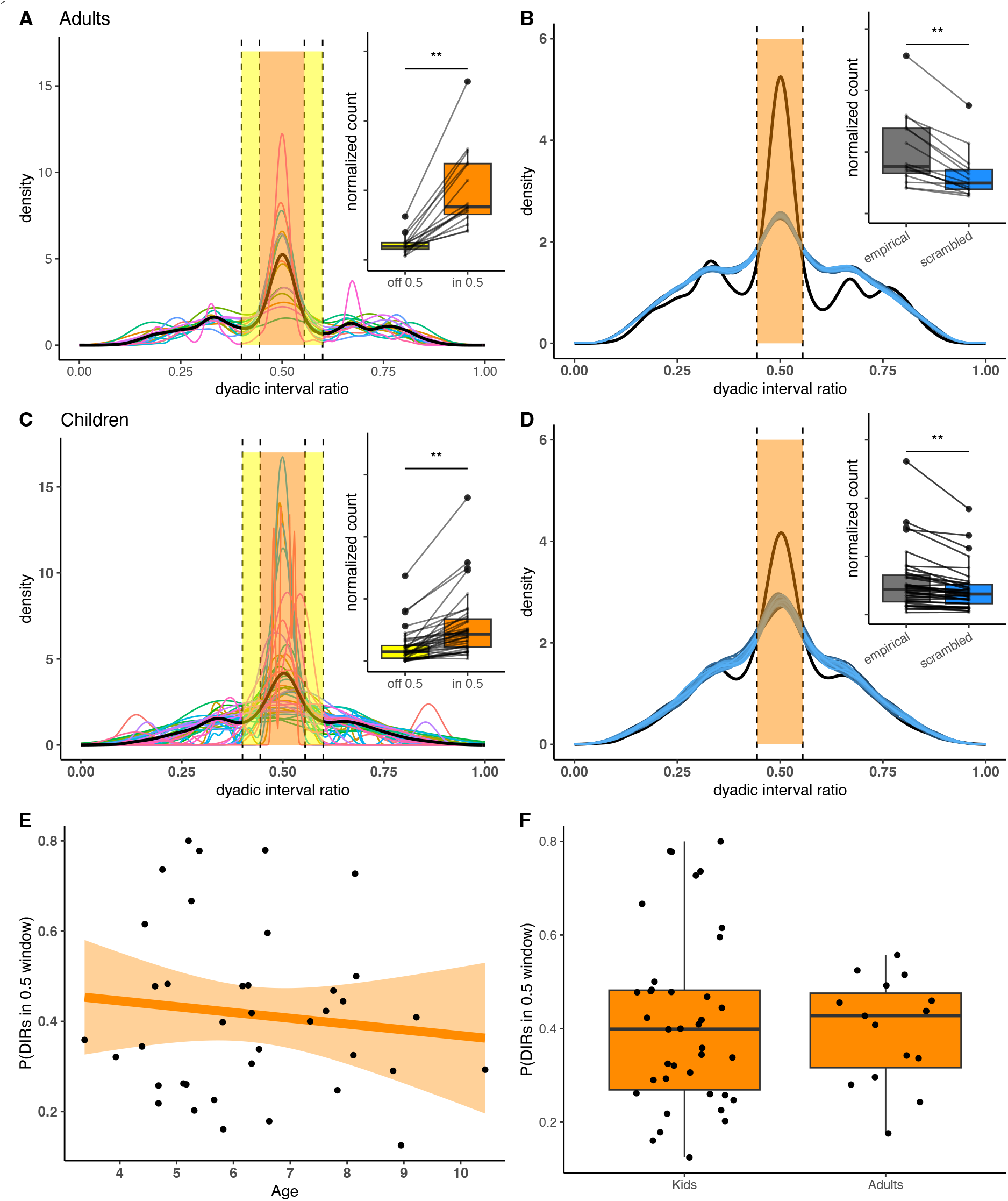
Non-musician adult and children’s improvised songs show isochrony. **Distributions of dyadic interval ratios of A** adult non-musicians’ songs show more ratios in the isochrony window (“on 0.5”) than outside of it (“off 0.5”) (each colored line is an individual participant’s distribution; the thicker black line is the population average). **B** Adults’ songs show more ratios inside this window than chance (each blue line plots the distribution of dyadic intervals in one of the 50 scrambled datasets; each point in the boxplot is a participant’s normalized count of ratios in the isochrony window, in the empirical data or averaged in the scrambled data). **C, D** Children’s improvised songs show the same pattern as adults. One child’s distribution of ratios is not plotted for clarity (the peak height would visually distort the rest of the graph; see Figure SI3). **E** There is no significant linear relationship between the children’s age and amount isochrony in their song, and F no significant difference between children and adul_6_ts in the amount of isochrony in their song. * p<0.05; ** p< 0.01

### 2. Children’s improvised songs also display isochrony

Similar to adults, children aged 3-10 years old spontaneously produce isochrony in their improvised songs (*Figure 2C* ; *t*(37) = 6.26, *p*< .001), at a higher-than-chance rate (*Figure 2D t*(37) = 4.488, *p* < .001). Additionally, there was no evidence for an influence of age on the degree of isochrony expressed: there was no linear relationship between children’s age and proportion of ratios that fell in the isochrony window (*Figure 2E* ; *β* = -0.0512, *SE*(*β*) = 0.0711, *z* = -0.721, *p* = 0.471) and no significant difference between children and adults in proportion of ratios that fell in the isochrony window (*Figure 2F* ; *β* = - 0.045, *SE*(*β*) = 0.144, *z*= -0.312, *p* = 0.755).

### 3. Zebra finch song is less isochronous than chance through the lifespan

In line with a previous report^10^, songs from our large colony sample of semi-naturalistically reared adult zebra finches (n = 61) showed significantly more dyadic interval ratios in the isochrony window than in the off window (*Figure 3A*; *t*(60) = 2.262, *p* = 0.027). While a significant difference in ratios in the on- and off-isochrony windows has previously been interpreted as the presence of isochrony in these vocalizations, further analysis reveals that the magnitude of this peak in the isochrony window is significantly lower than would be expected by chance (*Figure 3B*; *t*(60) =-6.557, *p* < .001).

**Fig. 3.**
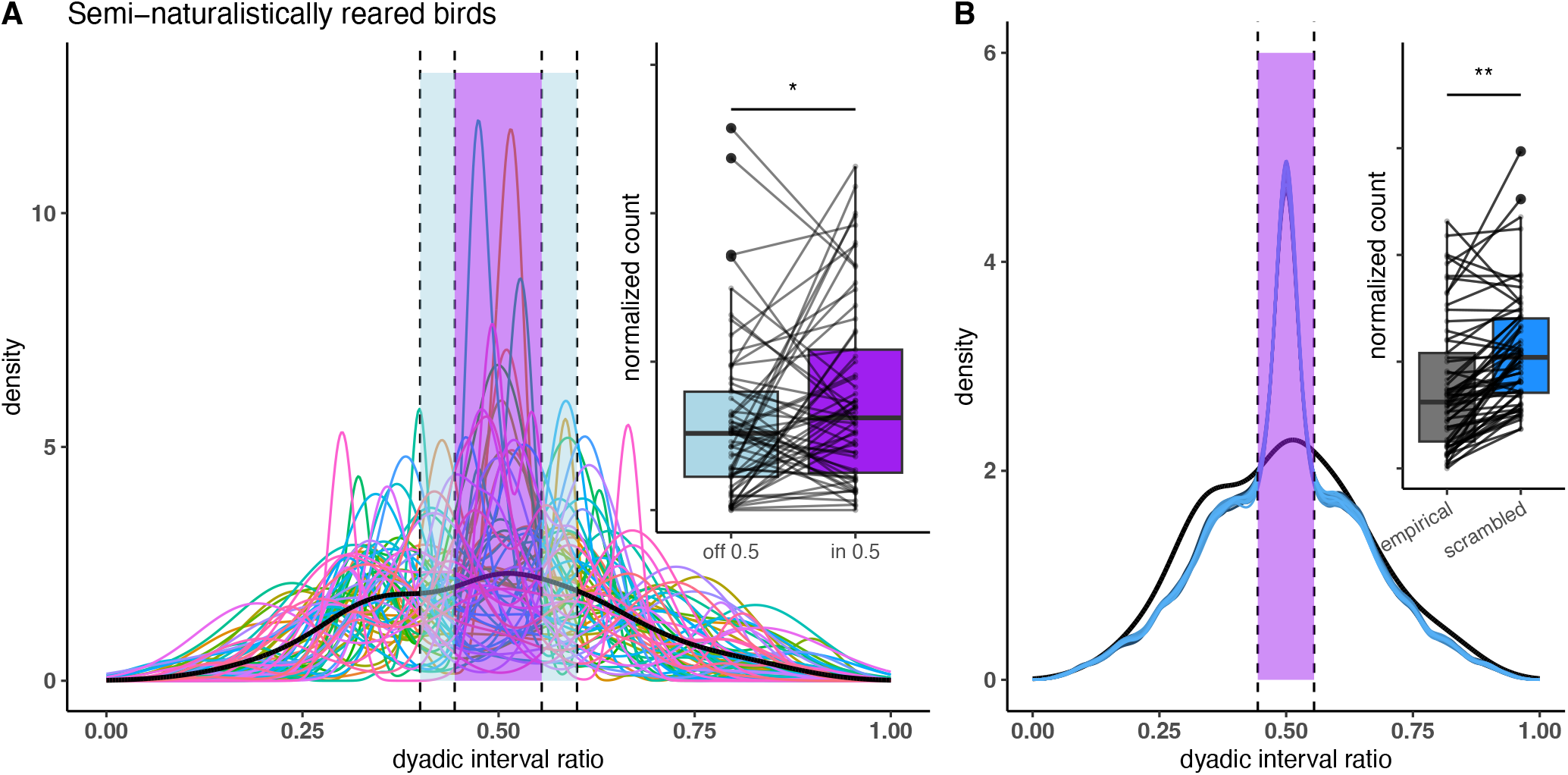
Zebra finch song shows a peak in ratios in the isochrony window that is lower than expected by chance. **A** Each colored line represents the distribution of dyadic interval ratios of an individual adult zebra finch. The thicker black line indicates the population average distribution of dyadic interval ratios. There are significantly more ratios within the isochrony window (“on 0.5”) than outside it (“off 0.5”), however **B** the height of this peak was significantly lower than expected by chance. The thicker black line is the same population average as in **A** and each blue line is the distribution of dyadic interval ratios as reconstructed from each of the 50 scrambles of these same birds’ data. * p<0.05; ** p< 0.01

In the longitudinally recorded birds, neither juvenile nor adult birds showed significantly more ratios in the isochrony window than in the off window (*Figure 4 A* and *B*; juveniles: *t*(15) = 1.781, *p* = 0.095; adults: *t*(15)= 2.013, *p* = 0.062). And similarly to the colony bird population, this incidence of ratios in the isochrony window was significantly lower than would be observed by chance for both juvenile (t(15) = -2.289, p = 0.037) and adult birds (t(15) = -2.944, p = 0.01; Figure 4). There was no effect of age on the incidence of isochrony production (β = -0.08, SE(β) = 0.215, z = -0.392, p = 0.695).

**Fig. 4.**
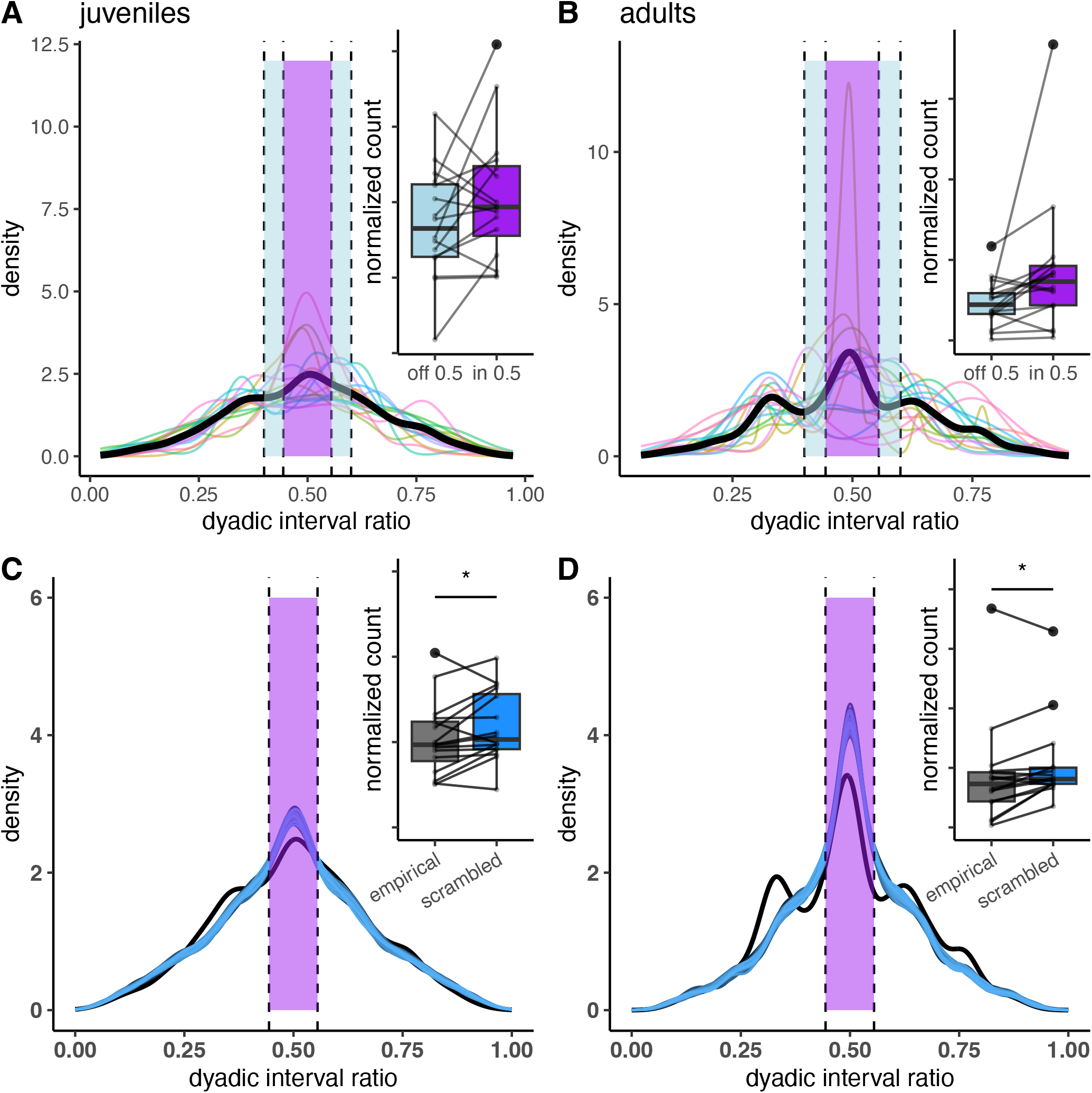
Zebra finch song is less isochronous than chance through the lifespan. Distributions of dyadic interval ratios from recordings of the same birds when they were **A** juveniles and **B** adults. Neither age group showed a significantly higher number of ratios in the isochrony window (“on 0.5”) than outside it (“off 0.5”). **C, D** Both juvenile and adult birds showed less isochrony than would be expected by chance. *\* p<0*.*05*

### 4. Zebra finches do not appear to learn isochrony naturally

We found no evidence for the learning of isochrony: the amount of isochrony (even if below chance) in the songs of tutor birds was not significantly predictive of the amount of isochrony in their pupils’ songs *Figure 5* ; *β* = 1.27, *SE*(*β*) = 0.9, *z* = 1.48 *p* = 0.138).

**Fig. 5.**
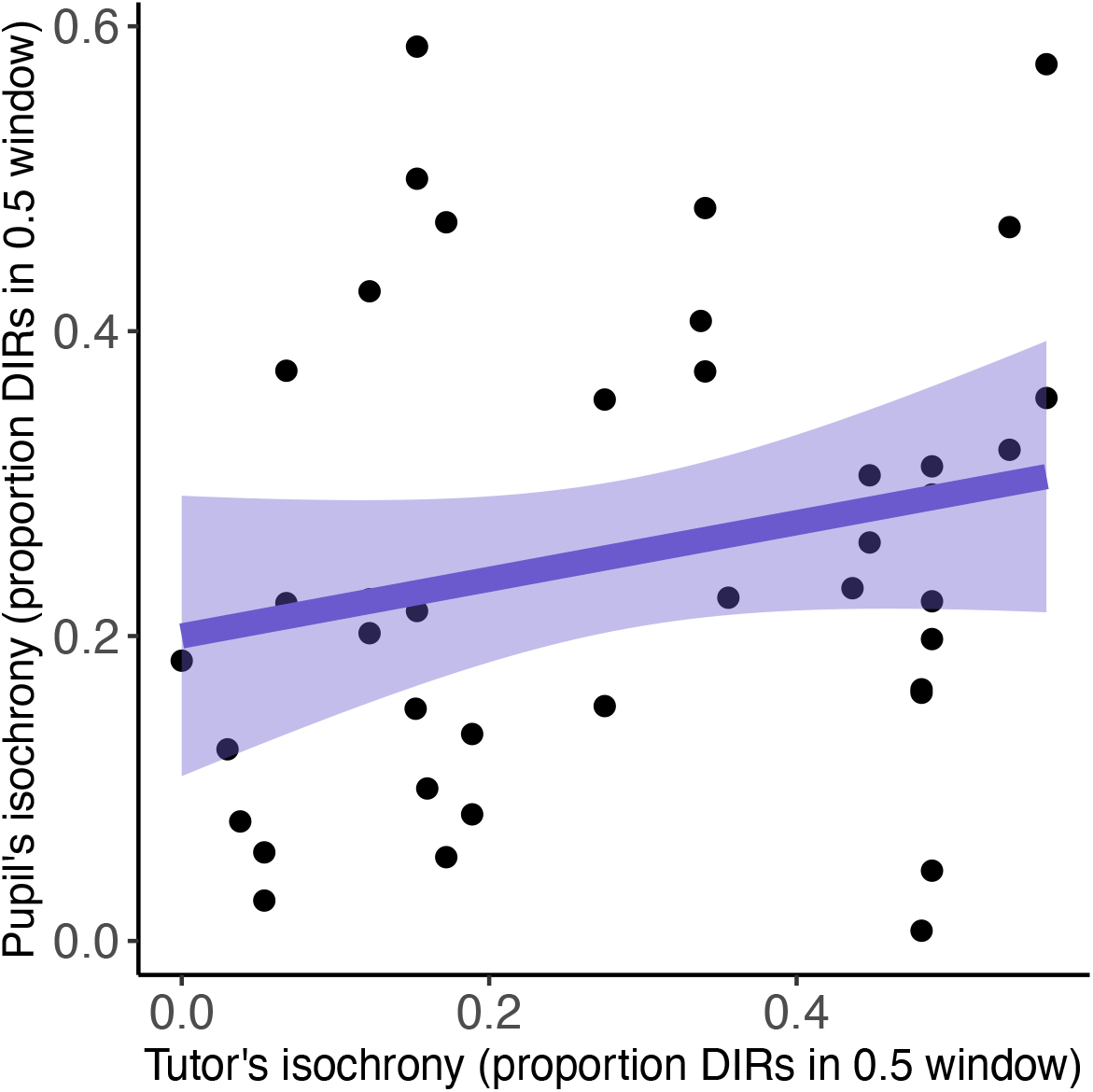
The lack of correlation of isochrony between a tutor and its pupil’s song suggests that zebra finches do not learn to produce this rhythm. The amount of isochrony in a tutor bird’s song is not significantly related to the amount of isochrony in thei pupil bird’s song. Each point is a tutor-pupil pair

## Discussion

We studied vocal rhythms in humans and zebra finches, using naturalistic and spontaneous expressions of vocalizations from each species. Despite that both species show early and socially-driven learning for the temporal features of their vocalizations, we find that only human song demonstrates isochrony to a degree greater than expected by chance, and that this species difference appears consistent through development (in the age ranges considered here). The present findings for improvised human song expand on the ecological validity of previous corpus^2^ and tapping^3^ studies that indicate how common isochrony is in human musicality. Our findings replicate a previous report of zebra finch song showing a high density of dyadic interval ratios around 0.5^10^. However, subsequent analyses indicated that the magnitude of this peak was significantly lower than what would be observed by chance, in both juvenile and adult birds’ songs: zebra finch songs do not display isochrony in a way that is analogous to human song.

What underlies this species difference is unknown. It is possible that humans show this strong propensity to produce isochrony due to their greater experience with this rhythm, but two of our findings weigh against this possibility. First, if more isochrony exposure in humans leads to more isochrony production, we would have observed a developmental increase in isochrony production: this was not the case in our data, with isochrony production being rather constant over the 3-10 year age range, and between children and adults. If there is an experience-dependence for the perception or production of isochrony, it might be acting only very early in development. Second, if isochrony production in both species were a function of exposure, we would have observed that the degree of isochrony in a pupil’s song would mirror that of their tutor. However, we saw no evidence that tutor birds’ individual levels of isochrony expression was predictive of their pupils’. This latter finding is surprising, given previous literature documenting the fine learning abilities of zebra finches. We suggest that this lack of correlation is a result of the birds’ prioritization of reproducing low-level temporal features, such as syllable structure and gap durations, over global rhythmic structures (as previously suggested in^45–47^). We observed instances in which the general sequencing of sounds was similar between tutors and pupils, but where pupils may merge of spilt syllables from their tutor’s song^48^, leading to an altering of inter-onset intervals and, thus, dyadic interval ratios. Any isochrony observed in zebra finch song might therefore be incidental to their reproduction of low-level temporal features, in contrast to human musical rhythms that are defined by relative durations for the consistent production of isochrony through a range of low-level temporal features (humans produce isochronous songs throughout a range of tempi: see SI Figure 4).

Future studies that replicate this method among denizens of musical cultures with appreciably different rhythmic structures would be needed to better characterize whether the present results of early arising and developmentally stable rates of spontaneous isochrony in song is in fact species typical of humans. The method used here - of recording children’s songs online as supervised by their parents - is a scalable alternative to traditional in-lab recording methods. This scalability, in addition to this method’s intrinsic requirement for minimal participant instruction, lends itself to facilitating cross-cultural study. If humans have no predisposed rhythm inclinations and isochrony production is purely experience-dependent, we might expect to see, for instance, that Malian children’s improvised songs would show a prevalence of isochronous and non-isochronous rhythms in accordance with their prevalence in jembe music^49^. Replications of this work would also reveal whether the lack of an age effect persists in other samples: we found no evidence for an increase in spontaneous isochrony production with age, which was surprising given that prior research on body movements to music or manual tapping to isochronous stimuli^19^ report that this rhythm production becomes more accurate through the age range of ∼2 to 7 years old. If replicated, the vocal production of isochrony would appear to be more precocious than previously expected.

In addition to the note/syllable level isochrony considered here, isochrony may occur at different time scales. For example, the Australian pied butcherbird’s song, while not isochronous at the level of syllables, does show isochrony at the level of phrases, with inter-phrase intervals being consistently of similar duration, on average ∼7.5 seconds^43^. In the present study, we employed a prevalent syllable-based segmentation for zebra finch song, where adjacent elements are delineated with brief silences. This is the same form of segmentation used in prior investigations of isochrony perception and production in birdsong^10,11^. Alternatively, one could also consider that syllables in bird song can consist of one or more “gestures”, where inter-gesture onsets have been reported to align with isochronous events more than would be expected by chance (see^33^, though this study also reports isochrony at the level of syllables, unlike our own data). Future work on the biological basis of rhythm in animals should first make explicit at what level (gestures, notes, phrases) we would expect to observe isochronous organization, in which species, and why. If the function of isochrony, as a perfectly predictable rhythm, is to serve inter-individual coordination and/or synchronization^50^, naturalistic interactions between individuals should show coordination at the levels of segmentation considered. Such an interpretation would be aligned with findings that isochrony is a defining feature of vocalizations that we chorus in, like song, and not a defining feature of turn-taking vocalizations like speech^38,51^ or zebra finch song, where these birds do not chorus, and deliberately adjust the timing of their singing to avoid overlapping with an interlocutor^45^.

The species difference in vocal isochrony we report here has broader implications about the biological basis of musicality. Humans produce isochrony in their improvised songs from a young age. Zebra finches do not produce this rhythm above chance, and despite their ability to learn many temporal features of song, isochrony does not seem to be one of them. Much work is ongoing to determine what necessary and sufficient conditions lead a species to exhibit the spontaneous production of this rhythm. One popularly evoked condition is the capacity for vocal learning^6,13,52,53^. The vocal learning hypothesis^34^ suggests that vocal learning neural circuitry is necessary for a species to express beat perception and synchronization, and many have tested whether vocal learner species also produce rhythmic vocalizations. Zebra finches learn their song during development but do not learn new vocalizations as adults, in contrast with humans and parrots. Despite their robust vocal learning abilities, evidence for isochrony in zebra finch song is weak. The present results therefore indicate that if vocal learning is in fact a driver for the *production* of learned, rhythmic patterns, it may be only the very high end of the vocal learning spectrum that produces this rhythm spontaneously in their learned vocalizations. This could be experimentally probed by investigating the spontaneous songs of parrots (e.g., cockatoos), currently the only non-human species conventionally considered very high on the vocal learner spectrum^34^. On the other hand, other frameworks propose that rhythm has its roots in coordinated group dance^36^ or vocal synchrony^38^, which, at time of writing, there is no evidence that parrots engage in; such frameworks may predict that parrots would not show spontaneous isochrony in their learned songs. Until such demonstrations, the trait of having an early-arising and minimally explicitly trained ability to produce spontaneously isochronous complex learned vocalizations appears to be unique to human song.

## Author contributions

Conception M.B., M.W., I.P., J.T.S.; experimental design and implementation M.W., J.T.S., D.S.; data processing M.B., M.W., D.S.; analysis and visualization M.B.; writing M.B. J.T.S., I.P.; editing M.B., M.W., I.P., J.T.S.

## Acknowledgements

This research was supported by the Fonds de recherche du Québec – Nature et Technologies (FRQ-NT) PR-299652 (J.T.S., I.P.); the Center for Research on Brain, Language, and Music (M.B., J.T.S.); a doctoral training grant from Fonds de Recherche du Québec Nature et Technologies (M.B. https://doi.org/10.69777/348674); and Natural Sciences and Engineering Research Council of Canada, Grant/Award Number: 05016 (J.T.S.) and 03113 (I.P.). We thank Daria-Salina Storch for gathering longitudinal bird recordings, Angela Wang and Logan James for gathering semi-naturalistically raised bird population recordings, Nicola Teolis for gathering children recordings, Alexandra Leblanc L’Ecuyer for segmenting children recordings. We also thank Ani Patel, John Iverson, and Jessica Grahn for important feedback and discussion.

## Notes

### Competing Interest Statement

The authors have declared no competing interest.

