## Supplementary material for "Humans, but not songbirds, produce spontaneously isochronous song through the lifespan": SI_tab1_fig1-4

### Supplemental Information

The improvised songs from human adults and children were consistent with previous studies on a number of features. Children's songs were significantly faster than adults' with smaller IOIs, m(children) = 477ms, m(adults) = 572ms, $t(36.7) = -3.3$, $p = 0.002$), consistent with children's faster spontaneous tempi in tapping experiments[54]. There was, however, no significant decrease in mean IOIs within the age range of 3-10 years old studied here (pearson correlation estimate between age in years and mean IOI $r(36) = -0.222$, $p = 0.18$). Adults' improvised lullabies were slower than their improvised dance songs (m(lullaby) = 651 ms; m(dance) = 405 ms, $t(14) = 0.246$, $p < .001$), consistent with previous cross-cultural studies of these song types[55,56]. These improvised songs therefore appear to corroborate the results of the study of tempo in other settings. Data availability differed per population, with there being more song from adults than children (see SI Table 1 ). For those unfamiliar with zebra finch vocalizations, we note that zebra finch song is much faster than human song, with much shorter inter-onset intervals (see SI Table 1 and SI Figure 2 for plots of distributions of IOIs for all populations). In humans, the adults' IOIs (mean 575ms) indicate that they were singing at a rate that was slower than previously reported spontaneous production rate when singing familiar songs of 330ms[57]. In order to assess timing variability, we computed the coefficient of variation (sd(IOI)/mean(IOI) as in58), and found no significant difference between children and adults (t-test between children's mean 0.6 and adults' mean 0.6 CVs, $t(49.6)= 0.096$, $p = 0.924$) or as a function of age among the children (spearman correlation between age and CV $t(36)= 0.4$, $p = 0.702$).

| Species | Age | Mean number of notes/syllables | SD number of notes/syllables | Mean IOI (ms) | SD IOI (ms) | CV |
|---|---|---|---|---|---|---|
| Human | Adults | 1404.7 | 650.1 | 574.6 | 359.0 | 0.6 |
| | Children | 155.1 | 116.3 | 490.4 | 357.7 | 0.7 |
| Bird | Adults | 372.2 | 158.9 | 186.7 | 136.1 | 0.7 |
| | Juveniles | 506.7 | 137.7 | 194.0 | 167.2 | 0.9 |

**SI Table 1 | Descriptive statistics**

The methodological decision of *what* to segment in any particular species' vocalizations is not trivial. We adopted the prevalent practice of segmenting birdsong into "syllables" - these are delineated by small breaths that the birds take, and so are visible on a spectrogram as delineated by small epochs of silence. This breath-group segmentation is conventional in the characterization of the temporal features of birdsong (*e.g.*[11,29]), and is the segmentation method that led to prior reports of there being isochrony in bird song and other animal vocalizations[10]. We also employ a conventional method for segmenting human vocalizations, which, in contrast, does *not* segment a sound wave solely on the basis of brief silences, but at the level of "notes"; these are segmented (in TONY software, for example) at the junctions of rapid pitch and amplitude changes (*Figure SI 1* ). We find that human song segmented using the same software we used for the processing of birdsong (MATLAB package `evsonganaly`), also showing isochrony in non-musicians' songs (more ratios in the 0.5 window than outside: $t(14)= 6.118$, $p< .001$, paired t-test).

One may therefore raise the question of whether the detection of isochrony in the vocalizations of either species may change if using the same segmentation method across species. Indeed, prior work has been done on segmenting birdsong into so-called *gestures* or *segments*, the units between rapid changes in frequency and amplitude[33,58]. One could also segment birdsong at the level of *bouts* (perhaps analogous to a "phrase" in human song), as has been done in the characterization of the temporal features of the Australian pied butcherbird's song[43]. In the present study we seek to extend on prior research that claims to detect human-like rhythms in birdsong[10,33] - prior research that segmented human song into *notes* and birdsong into *syllables* - we employed these same methods in order to ensure interpretability of our findings with respect to these prior claims.

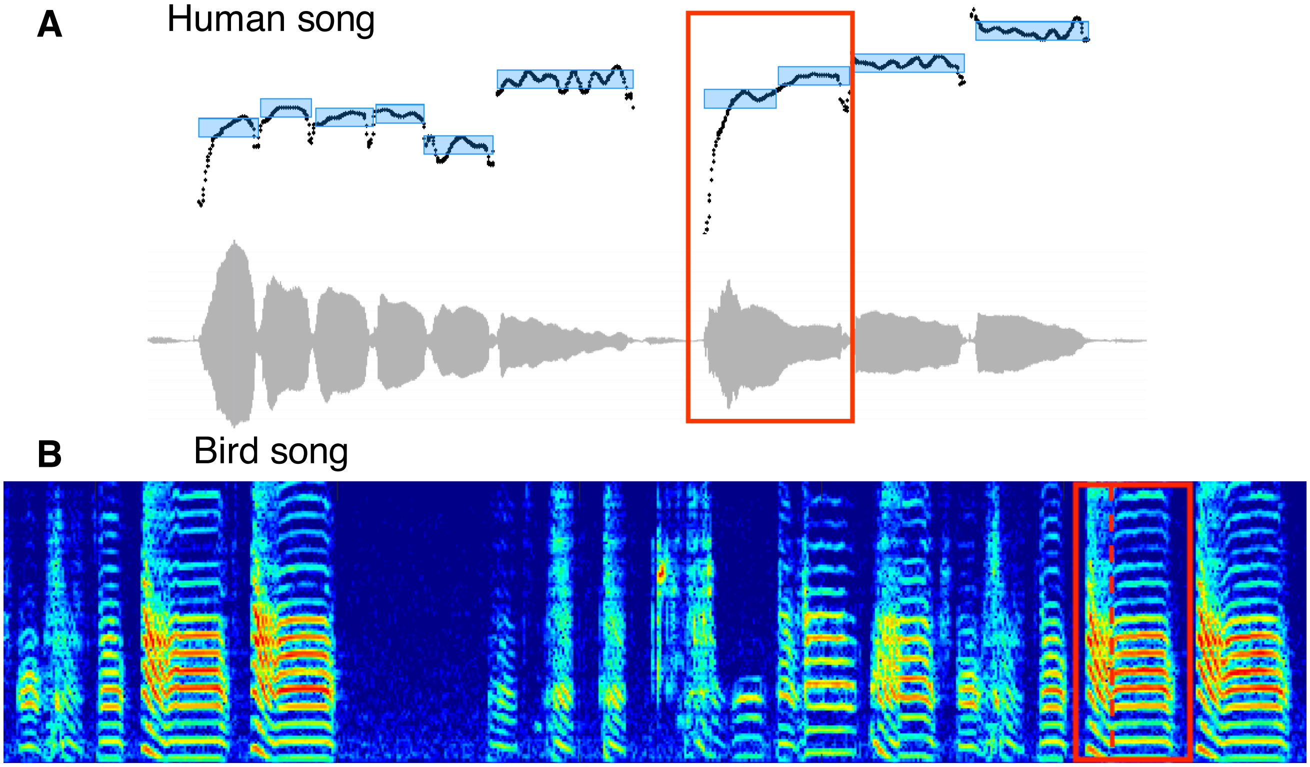


**Fig SI 1 | A** Human song is segmented into notes, at the joints of rapid transitions of pitch and amplitude. These notes are indicated by the blue rectangles. The red rectangle highlights a window where this signal is segmented into two notes, but this same signal, if it were to be segmented as birdsong conventionally is on the basis of only amplitude fluctuation, may have been segmented into one large syllable. **B** Birdsong is segmented into syllables. The red-highlighted region is conventionally segmented into a single syllable; if the same signal were to be treated as human song, it likely would have been segmented into two segments, along the transition in amplitude and frequency (marked with a vertical dotted line).

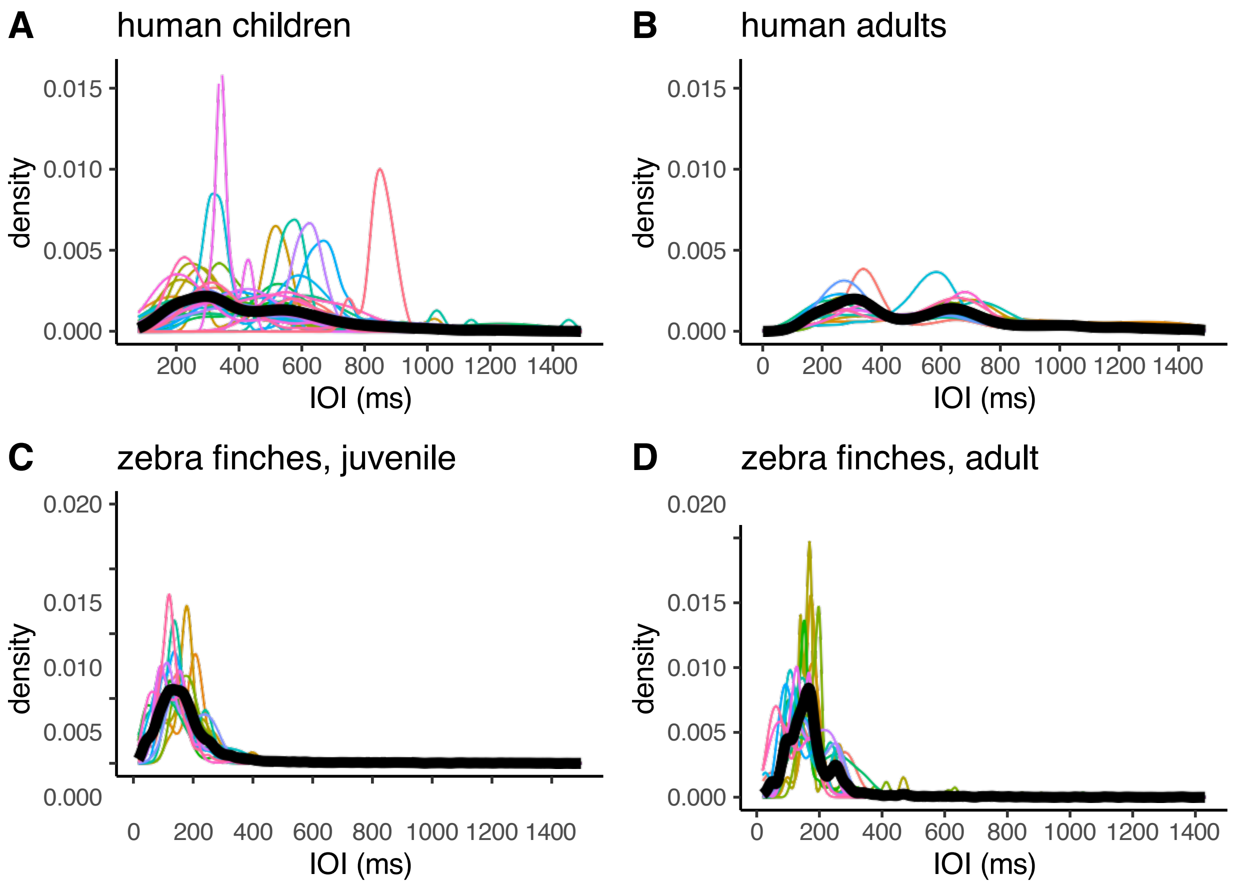


**Fig SI 2 |** Distributions of IOIs of spontaneous song from **A** children, **B** adults, **C** juvenile zebra finches, and **D** adult zebra finches (of the longitudinally-recorded population). For all panels, colored lines are individuals from the relevant population, and black lines are population averages.

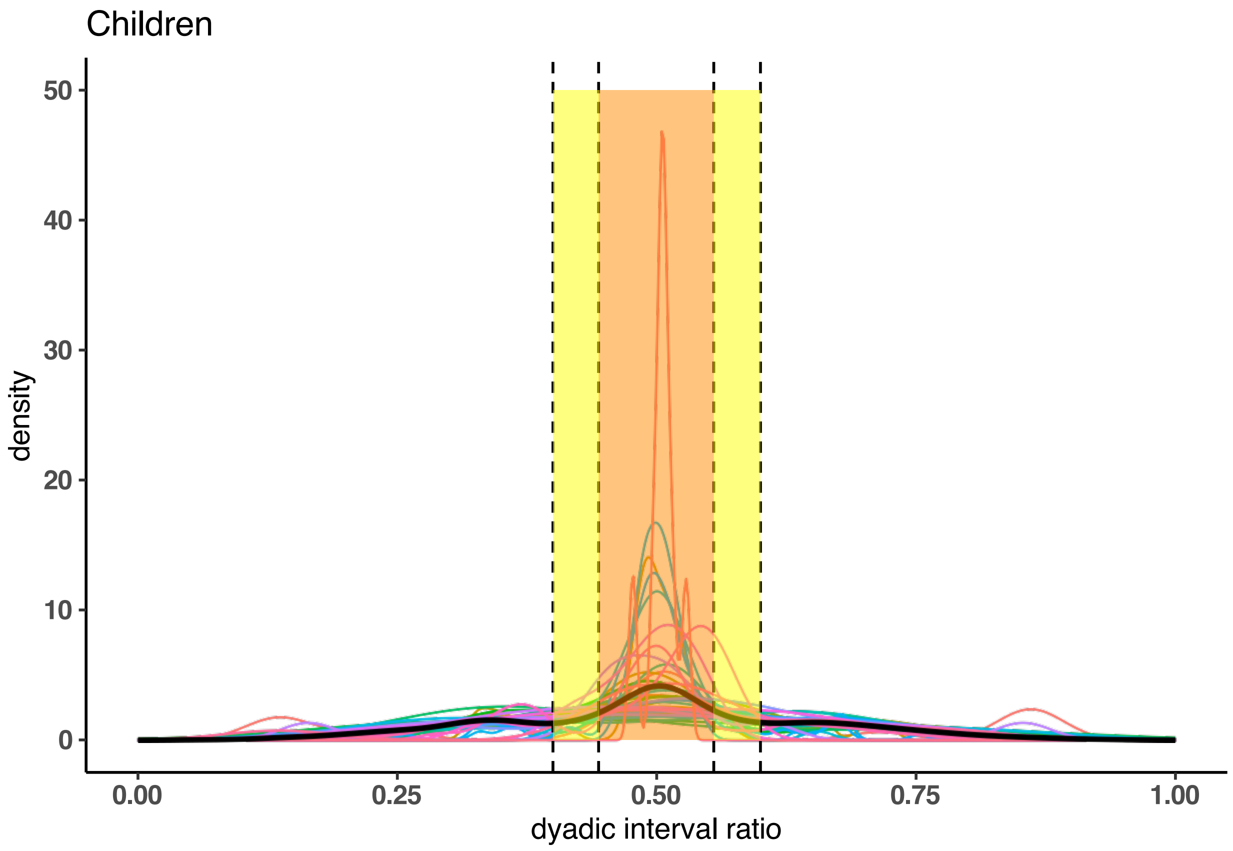


**Fig SI 3 |** Same as Figure 2A, plotting individual children's distributions of dyadic interval ratios, but with fulll y axis to accommodate one child whose DIR curve has a very narrow, high peak. Each colorful line is an individual child; the bolder, black line is the population average.

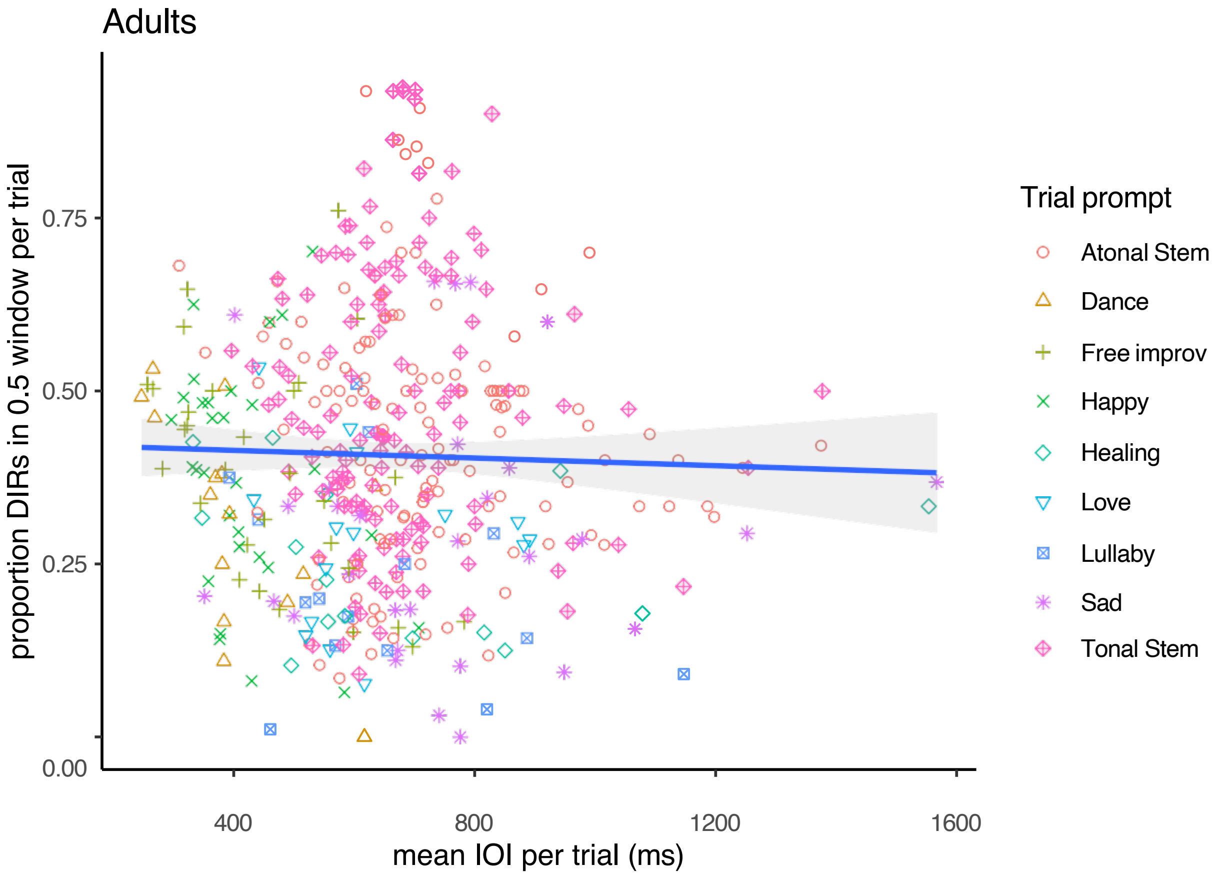


**Fig SI 4 |** Adults non-musicians' improvised songs showed no significant linear relationship between mean inter-onset interval and proportion of ratios in the isochrony window, indicating no evidence that adults are more inclined to produce more isochrony at certain tempi. We tested on effect of trial prompt on the basis of prior research (Trehub et al., 1993; Mehr et al., 2019) indicating that lullabies cross-culturally are slower than dance songs: we replicate this finding in our sample of non-musicians' improvised songs. [redacted] Different symbols indicate different song types, on the basis of prompt given for each trial.
